# Nanoplastics Penetration Across the Blood-Brain Barrier

**DOI:** 10.1101/2025.09.10.675462

**Authors:** Annemarie Ianos, Jessica Zhou, Tongxuan Qiao, Tao Wei, Baofu Qiao

**Affiliations:** Department of Natural Sciences, Baruch College, City University of New York, New York 10010, New York, United States; Syosset Senior High School, Syosset, New York 11791, New York, United States; Department of Chemical Engineering and Department of Biomedical Engineering, University of South Carolina, Columbia, 29208, South Carolina, United States

**Keywords:** blood-brain barrier, polymer, nanoparticle, passive permeation, free energy

## Abstract

Microplastics and nanoplastics (MNPs), originating from plastic degradation, have arisen to be a threat to ecology and human health. Alarmingly, the penetration of MNPs across the highly selective blood–brain barrier (BBB) poses an emerging and urgent risk, yet its molecular mechanism remains unexplored. In this work, using long-time-scale (over 27 μs) all-atom explicit solvent steered molecular dynamics, we examine the free energy of the passive permeation of four polymer nanoparticles: polyethylene, polypropylene, polystyrene, and polyethylene terephthalate. Polyethylene and polypropylene nanoparticles exhibited a remarkable preference for entering the BBB, attributed to their high hydrophobicity. Our study reveals that polymers can enter the BBB as polymerized nanoplastics and exit as dispersed polymer chains as the nanoparticles dissolve within the BBB. Further, the crystalline structure of polyethylene nanoparticles is found to adopt varying orientations. Our work advances the knowledge about the mechanism of nanoplastic penetration across the BBB, which could aid in the rational design of therapeutics for nanoplastic penetration inhibitors.

**TOC Graphics:** 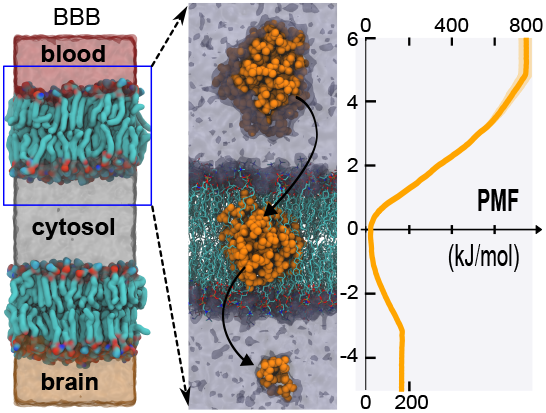

## INTRODUCTION

Despite their relatively short history of just over a century,^1^ synthetic polymers, such as plastics, have become integral to modern life. Their popularity stems from their unique properties, including durability, lightweight design, cost-effectiveness, and customizability. However, their durability, while advantageous in applications, poses a significant challenge when it comes to degradation after end-of-life use.^2^ For example, polyethylene terephthalate (PET) plastics can take approximately 450 years to degrade under ambient conditions.^3^ As a result, plastic pollution has become an emerging and urgent threat to the environment and human health, given the slow degradation and the generation of microplastics and nanoplastics (MNPs), which refer to plastic particles at the micrometer and nanometer scales, respectively. Human exposure to MNPs is ubiquitous: it is estimated that humans may ingest 0.1 - 5 g of microplastic particles weekly,^4^ or 39,000 – 52,000 microplastic particles annually.^5^ Specifically, the presence of MNPs has been reported in the food chain,^6-8^ human organs (*e*.*g*., liver^9^ and kidneys^10^), infant formula,^11^ and other sources.^12-14^ Recent studies have demonstrated that microplastics in the bloodstream can lead to neurobehavioral abnormalities,^15^ cellular toxicity in mammals,^16, 17^ and numerous diseases.^18, 19^

More alarmingly, MNPs have been very recently detected in the human brain,^20^ further highlighting the potential risks to human health. Specifically, through the autopsy of cadaver brains, livers, and kidneys, polyethylene (PE), followed by many other polymers, was found to be abundant in these organs, with the highest concentration in the blood-brain barrier (BBB) and brains of dementia patients. The BBB protects the human brain against foreign particles by strictly regulating the exchange of substances across the capillary walls. Its dysfunction is associated with neurodegenerative diseases, such as Alzheimer’s disease, Parkinson’s disease, and amyotrophic lateral sclerosis.^21^A study on mice found that exposure to polystyrene (PS) microplastics caused disruption in the BBB, cognitive defects, and inflammation of the hippocampus.^22^ These observations underscore the importance of analyzing the penetration of MNPs across human tissues, for instance, the BBB.

To the best of our knowledge, no molecular-level understanding has been reported on the MNPs crossing the BBB. In this study, we quantified the free energy of the passive permeation of polymer nanoparticles (NPs) crossing the BBB using long-time-scale all-atom explicit solvent steered molecular dynamics (sMD) simulations. Four types of small-sized polymer nanoplastics were examined: PE, polypropylene (PP), PS, and PET, which have a diameter of approximately 3.1 – 3.5 nm. These polymers are broadly employed in the polymer industry and everyday applications, including food packaging, wire insulation, carpets, appliances, automotive components, and many more. The passive permeation mechanism^23^ was found to dominate for PS nanoplastics with a diameter of 0.293 μm across the BBB,^24^ while the receptor-mediated transcytosis and the endocytosis mechanisms apply to micrometer-sized particles.^25^ It was found that PE and PP NPs displayed a remarkable preference for entering the BBB, followed by PS NPs and then by PET NPs. Our study reveals that the polymers enter the BBB as nanoparticles while exiting the BBB as single chains.

## RESULTS AND DISCUSSION

The BBB bilayer (**Figure 1A, B**) was generated using the CHARMM-GUI web server.^26^ The composition of the bilayer, representing the apical BBB bilayer,^27^ is listed in Supporting Information Table S1. It was equilibrated 0.8 μs using the CHARMM 36m potential. ^28^ The calculated cross-sectional area per lipid (APL) supports the accuracy of the simulation (Supporting Information Table S2). Hereafter, the BBB refers to the apical bilayer of the BBB.

**Figure 1.**
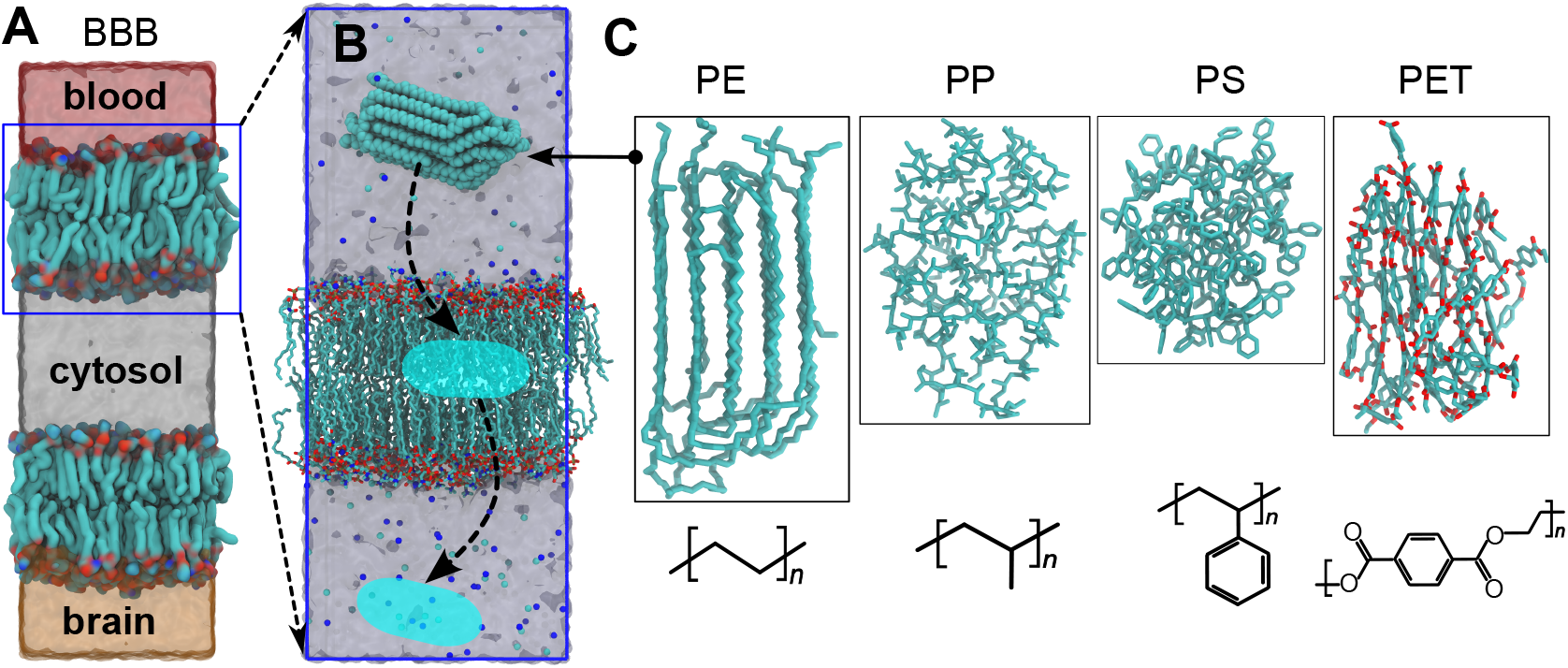
Schematic representation of (**A**) the BBB, and (**B**) a PE nanoparticle (colored in cyan) crossing the apical layer of the BBB. The Na^+^ and Cl^-^ ions are represented by the blue/cyan dots, respectively. (**C**) Four types of nanoplastics are employed, composed of PE, PP, PS, and PET. Each NP consists of 10 polymer chains and has a diameter of approximately 3.4/3.5/3.1/3.1 nm for the PE/PP/PS/PET NPs, respectively. The chemical formulas are presented below. The polymers have similar molecular weights (around 1 kDa) with the degree of polymerization *n* = 36/24/9/6 for PE/PP/PS/PET, respectively.

To mimic nanoplastics, we simulated four types of polymer NPs composed of PE, PP, PS, and PET, respectively (**Figure 1C**). Among these, PE microplastics have been found to be particularly prevalent in the brains of deceased individuals, followed by PP and several other polymers,^20^ underscoring the urgency of this research. Each NP was assembled with 10 polymer chains and had a diameter of approximately 3.1 – 3.5 nm, representing small-sized nanoplastics within the generally defined size range of 1 – 100 nm.^29, 30^ All polymer chains have similar molecular weights (around 1 kDa) to allow for comparison of their interactions with the BBB.

### Free energy of polymer nanoparticle insertion into the BBB

sMD simulations ^31^ were conducted to quantify the free energy of the polymer NP insertion into the BBB bilayer. For each system, long-time-scale (4.2 μs) all-atom explicit solvent sMD simulations were carried out using the pulling module of the GROMACS package.^32^ The potential of mean force (PMF) was then calculated using the Weighted Histogram Analysis Method (WHAM) algorithm,^33^ which is presented in **Figure 2**.

**Figure 2.**
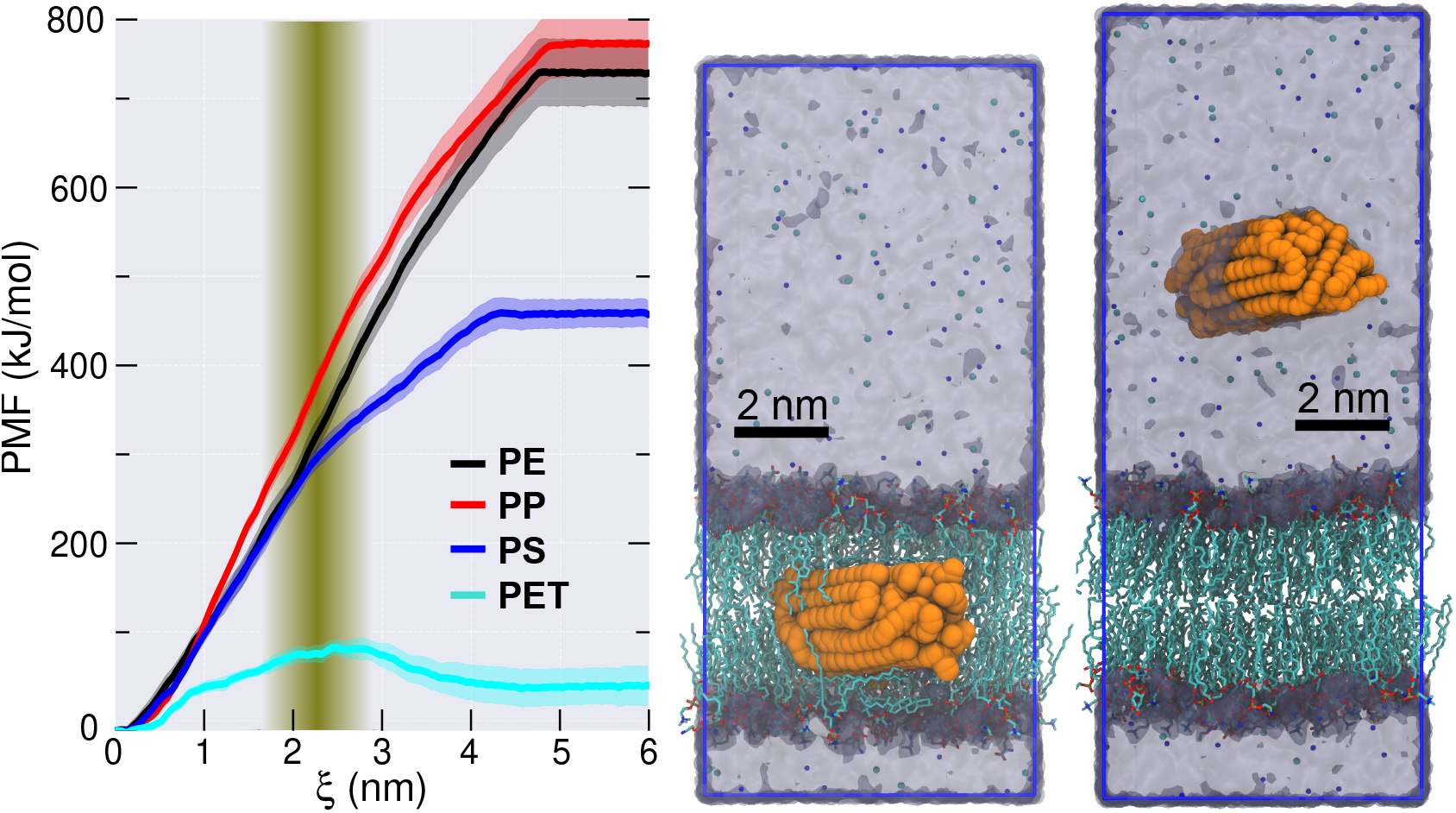
Free energy profile of polymer nanoparticle insertion into the BBB. The error bars represent the standard deviations of 4 blocks with 10 ns each. ξ stands for the Z-dimensional distance between the center-of-mass (COM) of the BBB and the COM of the polymer nanoparticles. The shadowed vertical region represents the distribution of the lipid phosphorus atoms in the control system (Supporting Information Figure S1), defining the interface of the BBB bilayer. Presented on the right are the representative simulation snapshots that show PE nanoparticles (in orange) located in the interior of the BBB (ξ = 0) and the water phase (ξ = 7 nm). The Na^+^ and Cl^-^ ions are represented by the blue/cyan dots, respectively. Simulation snapshots of the other systems are in Supporting Information Figure S2.

The free energy of the nanoparticle insertion into the BBB bilayer was obtained to be -739 ± 37 kJ/mol for PE, -772 ± 37 kJ/mol for PP, -469 ± 16 kJ/mol for PS, and -50 ± 22 kJ/mol for PET. The free energies align with the hydrophobicity of these polymers that PE ≈PP > PS > PET: The water contact angles on solid polymers have been experimentally reported to be 86° – 93° for PE, 85° – 96° for PP, 72° - 86° for PS, and 72° - 77° for PET.^34^ PE and PP are highly nonpolar, whereas the styrene group on PS provides an elevated polarity compared to PE and PP. It was also suggested that the interactions between styrene and cholesterol could enhance the attraction between lipid membranes and PS NPs, thereby promoting PS NP insertion.^24^ In contrast to PS, the introduction of the ester functional groups in PET further improves the polarity. Consequently, with the increase in the polarity from PE/PP, to PS, to PET, their hydration becomes more energetically favorable, and the interaction with the nonpolar interior of the BBB bilayer becomes less favorable, which collectively play a decisive role in the free energy of insertion in **Figure 2**.

### Anisotropic orientation of the PE nanoparticle

The PE chains assembled into a crystalline structure in water, similar to the solid crystalline phase of PE at room temperature.^35^ A strong orientation distribution was observed for the PE NP during its insertion into the BBB bilayer. To quantify its anisotropic orientation, we calculated the second-order Legendre polynormal orientational function.^36, 37^

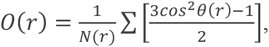

where *r* stands for the Z-dimensional distance between the BBB center and the center of the PE NP, *N*(*r*) denotes the number of PE NP neighbors at the distance *r* from the BBB center with a bin width of 0.1 nm, and *θ*(*r*) represents the angle between the normal of the BBB bilayer (the Z-dimension) and the principal vector of the NP. This orientation function has characteristic values: *O*(*r*) = –0.5 indicates that the nanoparticle is parallel to the bilayer plane (*i*.*e*., perpendicular to the bilayer normal), *O*(*r*) = 1 indicates that the nanoparticle is perpendicular to the bilayer plane (*i*.*e*., parallel to the bilayer normal), and *O*(*r*) = 0 denotes a random orientation.

The result is presented in **Figure 3**. Inside the BBB, the nanoparticle is found with O(*r*) = -0.464 ± 0.03, suggesting a strong parallel orientation relative to the bilayer plane (**Figure 3A**). This orientation is energetically favored as it increases the contact and hydrophobic interactions of the PE chains with the lipid tail groups. When the PE NP crosses the BBB/water interface, a strong vertical orientation (**Figure 3B, C**) dominates with a maximum O(*r*) = 0.968 ± 0.03 at *r* = 3.9 nm. In this orientation, the PE NP minimizes contact with the polar headgroups of the lipids. A similar orientation preference is observed for the “forever chemical”, per- and polyfluoroalkyl substances (PFAS) chains, at the lipid bilayer/water interface.^37^ In contrast, when the PE NP is in the water phase (more than 5 nm above the BBB center), it shows no preferred orientation with O(*r*) ≈ 0 (**Figure 3.C**). The random orientations displayed much larger error bars (0.0 ± 0.4) owing to a greater degree of freedom of 3-dimensional rotation in the water phase compared to that inside the bilayer and on the BBB/water interface.

**Figure 3.**
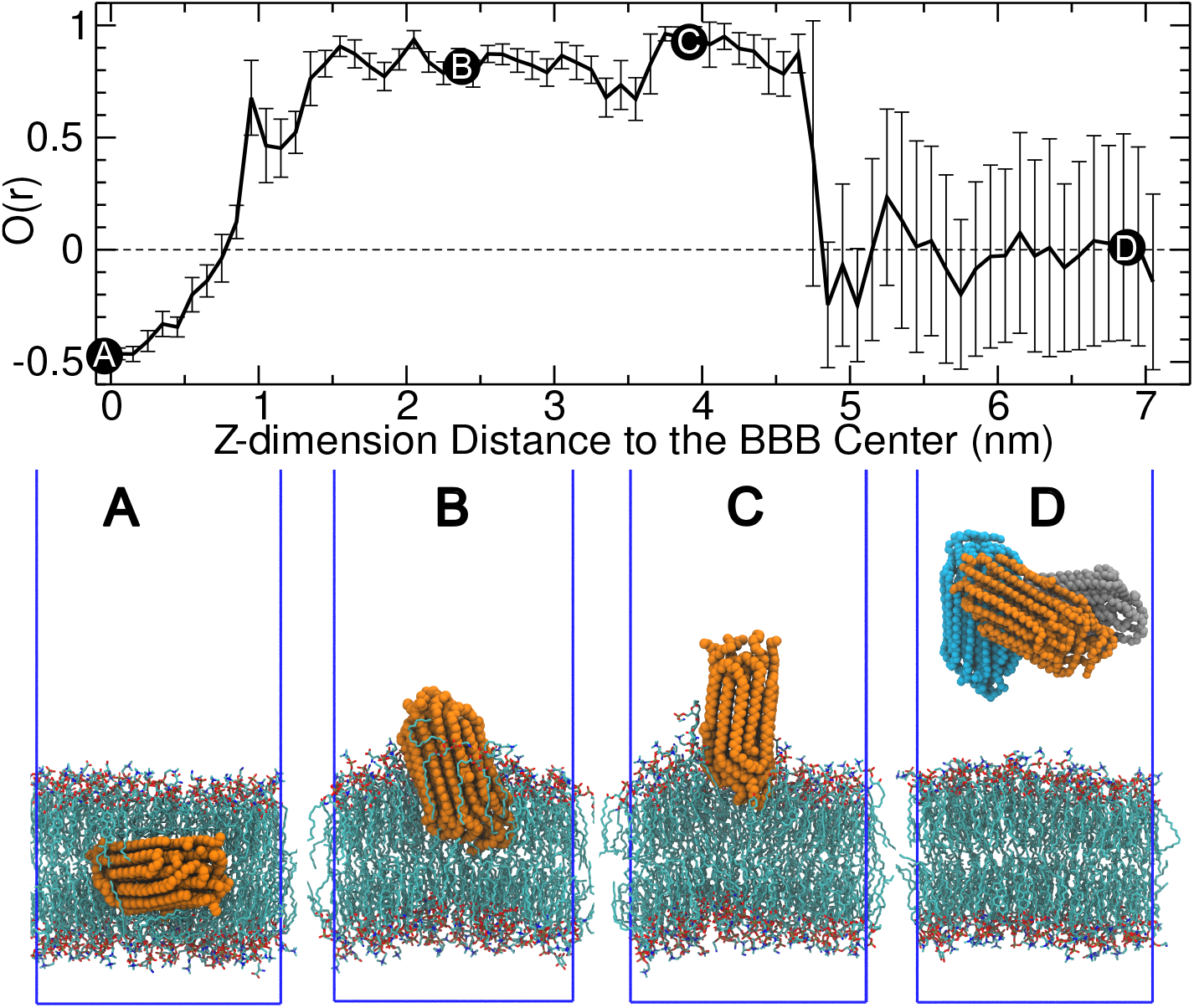
Orientation of the PE NP at varying Z-dimension height from the BBB center. The error bars represent the ensemble-based standard deviations. (**A**) In the BBB interior, the parallel orientation dominates between the membrane and the NP. (**B, C**) On the BBB/water interface, the vertical orientation dominates. (**D**) In the water phase, random orientations exist, with three different representative orientations (tilted in orange, vertical in cyan, and parallel in gray) overlaid on the same plot.

### Free energy of a single polymer chain exiting the BBB

The high free energies for the insertion of the PE, PP, and PS nanoparticles into the BBB (-772 ∼ -469 kJ/mol in **Figure 2)** indicate that entering the BBB bilayer is energetically favored, whereas the escape is highly challenging. A previous *in vivo* mice experiment suggested that BBB permeation is size-dependent and that green fluorescent signals were detected in mice brain tissues after 2 hours of exposure to small-sized PS NPs (*i*.*e*., 0.293 μm, where passive permeation dominates).^24^ This slow kinetics of BBB permeation is attributed to the fact that the free energy barriers are much higher than the driving force of the passive permeation mechanism, that is, fluctuations in the system’s potential energy (around ±100 kJ/mol in the systems with approximately 80,000 atoms investigated here).^38^

In a preliminary simulation, we observed that the polymer NPs could dissolve inside the BBB bilayer. Accordingly, we conducted classical MD simulations. Each simulation lasted 600 ns, which demonstrated the gradual dissolution of the polymer NPs when positioned in the BBB interior (**Figure 4A,B**), in agreement with a previous work using the MARTINI coarse-grained potential.^39^ Specifically, the dissolution was evidenced by the increases in the solvent-accessible surface area (SASA) of the polymer NPs and the potential energy between the BBB bilayer and the polymer NPs over the course of the 600 ns simulations. The dissolution is relatively more favored for the amorphous and nonpolar PP and PS NPs,^39^ in comparison with that of the PE (crystalline structure) and the PET (relatively polar) NPs.

**Figure 4.**
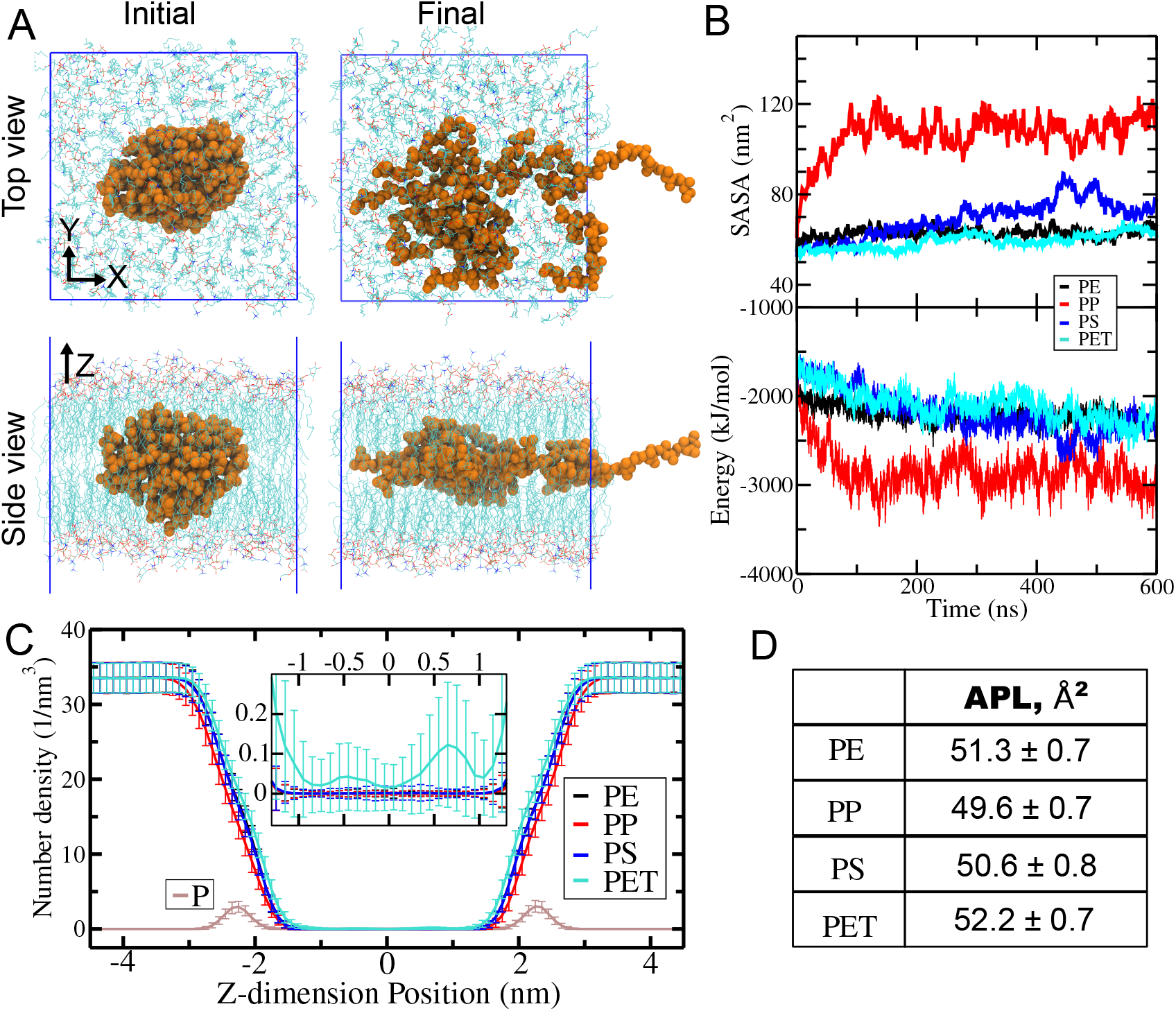
Dissolution of the NPs when embedded in the BBB interior. (**A**) Initial and final (600 ns) simulation snapshots of the PS nanoparticles. (**B**) The SASA of the polymer NPs and the total (coulombic + Lennard-Jones) interaction energy between the polymers and the BBB as a function of the simulation time. (**C**) Number density profile of water oxygen atoms in the presence of polymer chains dissolved in the BBB interior. The number density profile of the lipid phosphorus atoms (labelled as P) is included to indicate the BBB/water interface. The insert shows a close view of the central region, highlighting the elevated distribution of water in the presence of PET chains. (**D**) The APL with the polymer chains dissolved in the BBB interior, which are all larger than the value of 48.0 ± 0.7 Å^2^ in the control simulation without polymers.

Notably, when the PET chains are located at the BBB interior, the BBB interior becomes more hydrated (**Figure 4C**), similar to the observation for amphiphilic random heteropolymers when embedded in a lipid bilayer.^40^ Moreover, all BBB bilayers are found to expand when the NPs are positioned in the BBB interior, which is most pronounced for the PET NP (around an 8% increase; **Figure 4D** and Supporting Information Table S3). These findings suggest that the relatively polar PET polymers could deform the structure of the BBB bilayer, potentially facilitating BBB permeability.

Therefore, one plausible mechanism is that polymers enter the BBB membrane in their solid form (*i*.*e*., nanoplastics), dissolve in the hydrophobic interior of the membrane, and eventually exit the membrane as single polymer chains. To validate the escape of single-chain polymers, we conducted sMD simulations to quantify the free energy profiles of a single polymer chain (**Figure 5**). For all systems, the free energy barrier substantially drops for the single polymer chains compared to that for the polymer NPs: from 739 ± 37 to 145 ± 11 kJ/mol for PE, from 772 ± 37 to 153 ± 3 kJ/mol for PP, from 469 ± 16 to 97 ± 1 kJ/mol for PS, and from 50 ± 22 to 2 ± 8 kJ/mol for PET. Therefore, the penetration across the BBB is energetically much more favorable for single polymer chains.

**Figure 5.**
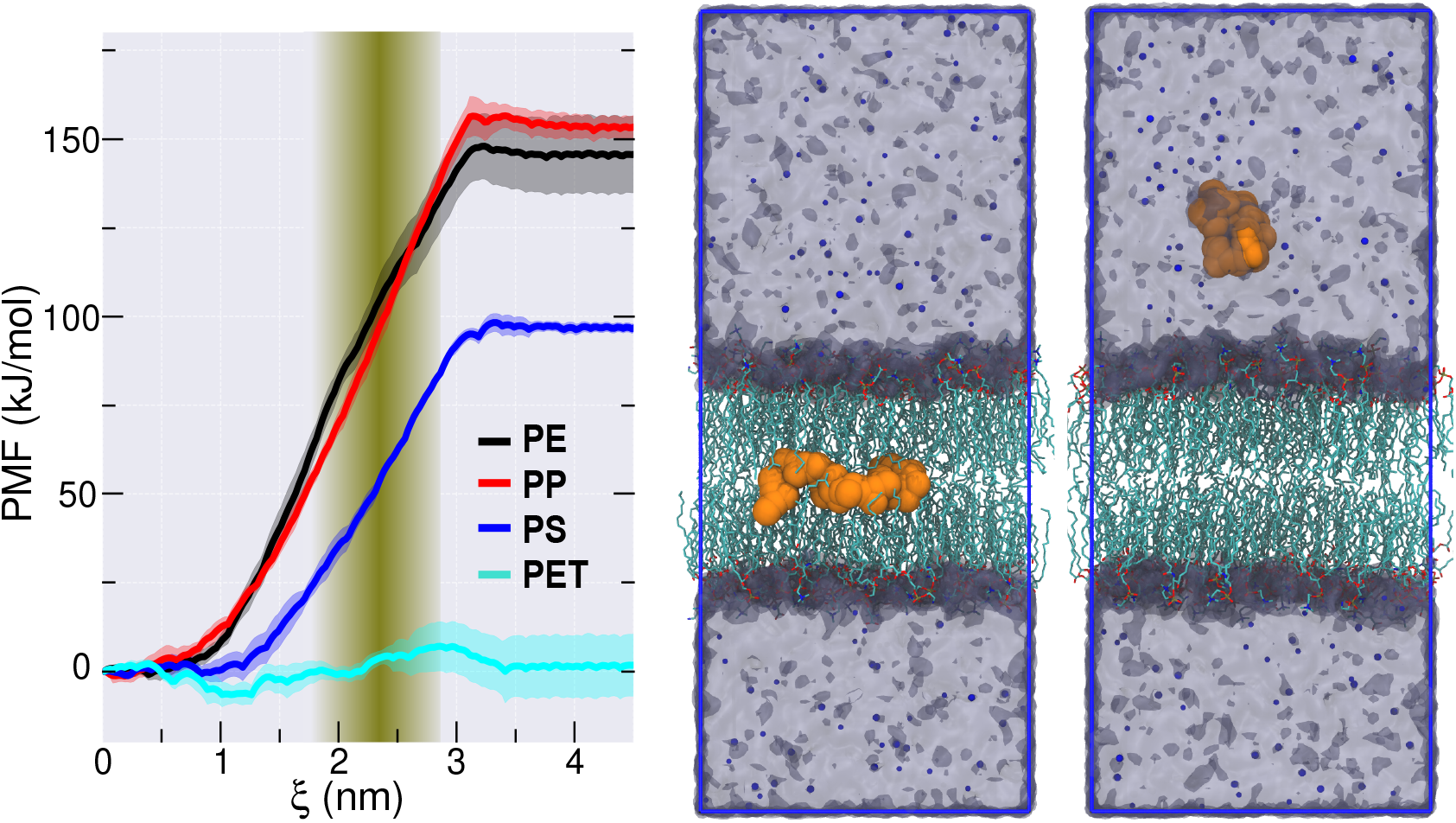
Free energy for a single polymer chain exiting the BBB bilayer. The shadowed vertical region denotes the interface of the BBB. Demonstrated on the right are the representative structures of the PP chain (in orange) located in the BBB interior (ξ = 0) and the water phase (ξ = 5 nm). The Na^+^ and Cl^-^ ions are represented by the blue/cyan dots, respectively. See Supporting Information Figure S4 for the simulation snapshots of the other systems.

Furthermore, the PMF profiles of the polymer NPs reach a plateau at a distance of 4 ∼ 4.7 nm (*e*.*g*., 4.7 nm for the PE NP) from the BBB center (**Figure 2**). Given that the radius of the NPs is around 1.55 - 1.75 nm (*e*.*g*., 1.75 nm for the PE NP) and the half of the thickness of the BBB bilayer is around 2.75 nm (Supporting Information Figure S1), it is supported that the polymer NPs display negligible long-range interactions with the BBB beyond their contact range. Accordingly, the PMF profiles for the single-chain polymers are converged at a distance of around 3.5 nm from the BBB center (**Figure 5**).

The overall free energy of the NP penetration across the BBB could be described as Δ*G*=Δ*G*(*NP entering BBB*) + Δ*G*(*NP dissolution*) + Δ*G*(*polymer chain exiting BBB*), where Δ*G*(*NP entering BBB*) denotes the free energy preference of a polymer NP entering the BBB (*e*.*g*., -739 ± 37 kJ/mol for PE) and Δ*G*(*polymer chain exiting BBB*) stands for the free energy barrier of a single-chain polymer escaping from the BBB (*e*.*g*., 145 ± 11 kJ/mol for PE). Although the absolute value of the free energy for polymer dissolution is computationally impractical owing to the slow dissolution kinetics, it could be reasonably assumed to be a negative value because of the spontaneous process of the polymer NP dissolution in the BBB interior. Taken together, it can be reasonably argued that the overall free energies for the entire penetration process Δ*G* are greater than -594/-619/-372/-48 kJ/mol for PE, PP, PS, and PET, respectively. Consequently, polymer penetration could be expected to be more energetically favorable under *in vivo* and *in vitro* conditions.

## CONCLUSIONS

We investigated the molecular mechanism and computed the free energy profiles for the passive permeation of PE, PP, PS, and PET NPs across the BBB using all-atom explicit-solvent sMD simulations. The energetic preference of the NPs entering the BBB follows the order PE (–739 ± 37 kJ/mol) ≈ PP (–772 ± 37 kJ/mol) > PS (–469 ± 16 kJ/mol) > PET (–50 ± 22 kJ/mol). These permeation free energies align well with the hydrophobicity of the polymers (PE ≈ PP > PS > PET): the high nonpolarity of PE and PP results in a strong preference for entering the BBB bilayer interior, while the elevated polarity of PET due to its ester groups leads to a substantial drop in the free energy preference for entering the BBB.

Non-biased atomistic MD simulations showed that these polymer NPs can dissolve when located in the BBB interior. This behavior is particularly pronounced for the nonpolar, amorphous PP and PS NPs, as evidenced by their calculated solvent exposure and interactions with the BBB. The high free energy barrier for polymer NPs exiting the BBB, combined with their dissolution in the BBB interior, suggests that polymers enter the BBB as polymerized nanoplastics but exit as single polymer chains. This is supported by the substantial drop in the free energy barrier for single polymer chains exiting the BBB.

We further found that the PE NP adopts different anisotropic orientations depending on its location: parallel to the bilayer plane when located in the BBB interior, vertical at the BBB/water interface, and randomly oriented when positioned in the bulk water phase. These orientations are believed to facilitate the BBB penetration. In contrast, the relatively more polar PET NP increases hydration of the hydrophobic BBB interior and expands the BBB bilayer, despite showing the least preference for entering the BBB.

Our study represents a first step toward understanding the molecular mechanism by which PE, PP, PS, and PET nanoplastics passively penetrate across the apical bilayer of the BBB. For a complete picture of the MNP penetration, further studies are needed on the factors, such as the nanoparticle size, the degree of polymerization of the polymers, polymer cross-linking, and other possible penetration mechanisms (*e*.*g*., transcytosis and endocytosis). These studies will aid in the rational design of therapeutics to prevent MNPs from crossing the BBB.^35, 41-43^

## Supporting information

Simulaton methods, Supplementary Tables and Figures.

## ASSOCIATED CONTENT

### Supporting Information

The Supporting Information is available free of charge at https://pubs.acs.org/doi/10.1021/XXX.

Methods. Supporting tables of the compositions of the BBB, the area per lipid. Supporting figures of BBB membrane thickness, the initial and final structure of the sMD, and the convergence of the PMF calculations.

## AUTHOR INFORMATION

### Author

Annemarie Ianos − Department of Natural Sciences, Baruch College, City University of New York, New York 10010, New York, United States

Jessica Zhou − Department of Natural Sciences, Baruch College, City University of New York, New York 10010, New York, United States

Tongxuan Qiao – Syosset Senior High School, Syosset, New York 11791, New York, United States

Tao Wei - Department of Chemical Engineering and Department of Biomedical Engineering, University of South Carolina, Columbia, 29208, South Carolina, United States

†: A.I. and J.Z. contributed equally.

### Author Contributions

A.I., J.Z., and T.Q. conducted the simulation, analyzed the data, and wrote the manuscript. T.W. contributed to the discussion and wrote the manuscript. B.Q. designed the project, conducted the simulation, analyzed the data, and wrote the manuscript.

### Notes

The authors declare no competing financial interest.

## ACKNOWLEDGMENTS

This work was supported by the Eugene M. Lang Foundation to B.Q.

