## Supplementary material for "Nanoplastics Penetration Across the Blood-Brain Barrier": Simulaton methods, Supplementary Tables and Figures.

### Table of Contents

**I. Methods ........ S2**

**II. Supporting Tables ........ S5**

**III. Supporting Figures ........ S7**

**References ........ S12**

### I. Methods

All-atom explicit solvent MD simulations were carried out to examine the polymer nanoparticles, the BBB, and their interactions. The open-source package GROMACS (version 2024.5) was employed.[1] The CHARMM 36m force field,[2] which is known to accurately describe lipid bilayers,[3] was applied to the lipids and metal ions ($Na^+$ and $Cl^-$). The CGenFF (CHARMM general force field) potential (version 4.6)[4, 5] was generated for the polymers using the CGenFF web server (cgenff.com). The recommended CHARMM TIP3P water model [6] was used with the structures constrained using the SETTLE algorithm.[7]

Unless otherwise stated, in all simulations, the neighbor searching was calculated up to a cutoff distance of 12 Å via the Verlet particle-based algorithm and was updated every 20 timesteps. The short-range Coulomb interactions were truncated at the cutoff distance of 12 Å, with the long-range interactions calculated using the Smooth Particle Mesh Ewald algorithm.[8, 9] The Lennard-Jones 12-6 interactions were switched off from 10 to 12 Å via the force-switch method. The temperature coupling was used via the V-rescale algorithm with the temperatures of the BBB, the polymers, and water/ions separately coupled at 303 K with a characteristic time of 1 ps. The semi-isotropic pressure was managed using the C-rescale algorithm at the reference pressure of 1 bar with a compressibility of $4.5 \times 10^{-5}$ $bar^{-1}$ and a coupling constant of 5.0 ps. The integration time step of 2 fs was used with all hydrogen-involved covalent bonds constrained using the LINCS algorithm.[10, 11]

**BBB bilayer preparation.** The BBB bilayer was generated using the CHARMM-GUI web server.[3] The composition of the BBB bilayer, representing the apical bilayer of the BBB,[12] is listed in Supporting Information Table S1. Here, a total of 192 lipids (including cholesterol) were used in the symmetric bilayer, with a water-to-lipid number ratio of 100 : 1. A control simulation of the BBB bilayer (without polymer NPs) with 150 mM NaCl was conducted for 0.8 μs. The equilibrated system has a size of 6.75 nm × 6.75 nm × 10.75 nm in X × Y × Z dimensions, respectively. We examined the cross-sectional area per lipid (APL) of the BBB at varying temperatures (Supporting Information Table S2). The calculated APL supported the convergence of the simulation within the first 0.05 μs. The APL was obtained to be 48.0 ± 0.7 $Å^2$ at 303 K, in agreement with prior simulation data of 45.3 $Å^2$ at 310 K.[12] The BBB bilayer thickness was estimated to be approximately 4.5 nm based on the Z-dimensional distributions of the lipid phosphorus atoms (Supporting Information Figure S1).

**Assembly of polymer nanoparticles.** Four types of polymer nanoparticles, mimicking nanoplastics, were simulated. Each nanoparticle was assembled with 10 polymer chains. The degree of polymerization is 36, 24, 9, and 6 for PE ($C_{72}H_{146}$, molecular weights (m.w.) = 1012.9 Da), PP ($C_{72}H_{146}$, m.w. = 1012.9 Da), PS ($C_{72}H_{74}$, m.w. = 939.4 Da), and PET ($C_{60}H_{50}O_{24}$, m.w. = 1155.0 Da), respectively. The molecular weights are similar for all polymer chains for the convenience of comparison. The polymer chains were generated using the CHARMM-GUI web server. The atactic configuration was employed for the PP and PS chains. Ten extended polymer chains were randomly placed in a simulation box with an edge length of 10 nm in vacuum and were then relaxed for 200 ns via the annealing process (gradually dropping the temperature from 500 K to 200 K). All ten polymer chains were condensed into a nanoparticle, which was subsequently dissolved in a cubic water box with an edge length of 6.6 nm. It was further equilibrated for another 200 ns with the isotropic pressure coupling. The obtained nanoparticles are presented in **Figure 1C**. Their diameters are estimated to be around 3.1 – 3.5 nm. The sizes are comparable to a recent work using the MARTINI coarse-grained potential, where PS nanoparticles were simulated interacting with a DOPC bilayer.[13] Notably, the PE chains form a crystalline structure, consistent with the crystalline solid phase at room temperature.[14] This further supports the accuracy of the CHARMM36m potential.

Furthermore, we calculated the densities of these nanoparticles. The estimated densities are 0.85 g/cm$^3$, 0.75 g/cm$^3$, 1.00 g/cm$^3$, and 1.28 g/cm$^3$ for the PE, PP, PS, and PET NPs, respectively. These values are slightly lower than the experimental data of 0.917-0.94 g/cm$^3$ for low-density PE,[15] 0.85 g/cm$^3$ for amorphous PP,[16] 1.04-1.05 g/cm$^3$ for PS,[17] and 1.38 g/cm$^3$ for PET.[18] The relatively lower densities obtained here are ascribed to the absence of high pressure when preparing these structures[14] and their lower molecular weights (approximately 1 kDa *vs.* tens of kDa or higher in practical applications).

**sMD simulations of nanoparticles**. The pre-equilibrated BBB and nanoparticle systems were merged to prepare the BBB/nanoparticle systems with a salt concentration of 150 mM. A classical (non-biased) MD simulation of 2 μs each supported the absence of the insertion of these NPs into the BBB bilayer, necessitating the sMD simulations.

Consistent with a previous work using the MARTINI coarse-grained potential,[19] we found that the polymer chains dissolve when embedded in the interior of the BBB (**Figure 4**). Their slow dissolution kinetics make it impractical for atomistic simulations to sample the full relaxation of the polymer chains in the sMD simulations. As a result, an elastic network [20] was introduced to preserve the solid-like structure of the polymer NPs. Specifically, a harmonic potential (bond type = 6 in GROMACS and force constant = 10,000 kJ/mol/nm$^2$) was applied to the alpha-C backbone atoms within the distance range of 3.0 – 9.0 Å.

The polymer NP was first pulled to the BBB center and equilibrated for 500 ns at 303 K. The insertion of the polymer NPs in the BBB interior was found to increase the APL by around 5% ~ 10% (Supporting Information Table S3). The sMD simulation was then conducted using the pulling code in GROMACS. It is a two-step protocol. Firstly, the COM of the polymer nanoparticle was pulled upwards along the Z-dimension from the BBB center to a distance of 7 nm, where a pulling force constant of 5000 kJ/mol/nm$^2$ was employed. To better preserve the structure of the BBB bilayer, a very slow pulling rate of 0.1 nm/ns was used. To prevent the flip of the lipid chains, position restraints were applied on their headgroups and tail groups. The simulation trajectory was saved every 10 ps. The distance between the COM of the nanoparticle and the COM of the BBB was calculated as a function of the simulation time.

Subsequently, the simulation configurations were extracted for a 1 Å interval of the COM-COM distance in the range of 0 – 7 nm, generating 71 windows. Umbrella sampling was carried out by setting the pull rate to zero, along with the removal of the position restraints on the lipids. Each sampling window was equilibrated for 60 ns, with a saving frequency of 100 ps per frame. Calculations of the APL supported the convergence of the simulations within the first 20 ns (Supporting Information Table S3). Therefore, only the last 40 ns were employed for data analysis. Around 4.26 μs of sMD simulation was conducted for each of the four systems, totalling around 17 μs. The final structures of the last windows are presented in Supporting Information Figure S2.

The potential of mean force (PMF) was calculated using the *gmx wham* program. The convergence of the PMF calculations is presented in Supporting Information Figure S3. To calculate the standard deviations of the free energy, the 40 ns simulation was divided into 4 blocks, with 10 ns each. The anisotropic orientation of the PE nanoparticle was calculated using an in-house script based on our previous work.[21]

**sMD simulations of single polymer chains**. These simulations are similar to those for nanoparticles above. The initial structures were prepared by deleting 9 out of the 10 polymer chains and equilibrated for 40 ns. For the sMD simulation, each sampling window runs for 50 ns, with the last 30 ns employed for PMF calculations. The simulations were conducted for the COM–COM distance range of 0 – 5 nm, which lasted around 10 μs in total. The convergence of the calculated PMFs is presented in Supporting Information Figure S4. To calculate the standard deviations of the PMFs, each 30 ns simulation was divided into 3 blocks, with 10 ns each. The configurations with the polymer chain located in the BBB center and the water phase are presented in Supporting Information Figure S5.

**Non-biased simulations on NP dissolution in the BBB interior.** We carried out classical MD simulations to examine the dissolution of the NPs when positioned in the interior of the BBB, where the constraints of the elastic network were removed to allow the full relaxation of the polymer chains. The simulation protocol was the same as that in the sMD simulations above, except that the pulling code was turned off here. Each simulation run for 600 ns, and the trajectory was saved every 1 ns. The dissolution of the polymer nanoparticles was characterized using the solvent-accessible surface area (SASA) and the interaction energy between the polymer chains and the BBB bilayer. The gradual convergence of these properties supported the slow dissolution kinetics of the polymers.

### II. Supporting Tables

**Table S1**. Composition in each leaflet of the BBB apical bilayer. [a]

| Type | POPC | CHOL | OSM | SLPC | SOPE | SAPE | SAPC | SAPS | SAPI |
|---|---|---|---|---|---|---|---|---|---|
| Number | 4 | 28 | 18 | 8 | 6 | 14 | 8 | 8 | 2 |
| Percent, % | 4.2 | 29.2 | 18.8 | 8.3 | 6.2 | 14.6 | 8.3 | 8.3 | 2.1 |

[a] POPC = 1-palmitoyl-2-oleoyl-sn-glycero-3-phosphocholine,

CHOL = cholesterol,

OSM = N-oleoyl-d-erythro-sphingosylphosphorylcholine,

SLPC = 1-stearoyl-2-linoleoyl-sn-glycero-3-phosphocholine,

SOPE = 1-stearoyl-2-oleoyl-sn-glycero-3-phosphoethanolamine,

SAPE = 1-stearoyl-2-arachidonoyl-sn-glycero-3-phosphoethanolamine,

SAPC = 1-stearoyl-2-arachidonoyl-sn-glycero-3-phosphocholine,

SAPS = 1-stearoyl-2-arachidonoyl-sn-glycero-3-phospho-L-serine,

SAPI = 1-stearoyl-2-arachidonoyl-sn-glycero-3-phosphoinositol.

**Table S2.** Area per lipid (APL, $Å^2$) at varying temperatures in the control system (*i.e.*, without nanoparticles).

| **BBB apical bilayer** | | | | | |
|---|---|---|---|---|---|
| **Temperature** | **303 K** | **350 K** | **400 K** | **450 K** | **500 K** |
| APL (This work) | 48.0 ± 0.7 | 53.5 ± 0.8 | 60.0 ± 1.2 | 67.5 ± 1.5 | 75.2 ± 3.3 |
| APL [a] | 45.3 @310 K | 52.3 | 60.6 | 67.1 @440 K<br>70.3 @460 K | 73.9 |

**Table S3.** Area per lipid (APL, /$Å^2$) in different simulation conditions in the presence of nanoparticles.

| | | **BBB_PE** | **BBB_PP** | **BBB_PS** | **BBB_PET** | **Control** |
|---|---|---|---|---|---|---|
| Interior [a] | Steered MD | 51.7 ± 0.5 | 51.1 ± 0.5 | 50.2 ± 0.6 | 53.6 ± 0.6 | - |
| | unbiased MD | 51.6 ± 0.6 | 50.5 ± 0.6 | 50.2 ± 0.7 | 52.8 ± 0.6 | |
| Outside [a] | Steered MD | 47.9 ± 0.5 | 47.4 ± 0.6 | 47.4 ± 0.5 | 48.0 ± 0.9 | 48.0 ± 0.7 [b] |
| | unbiased MD | 47.9 ± 0.8 | 47.9 ± 0.6 | 47.8 ± 0.7 | 47.8 ± 0.7 | |

a) "Interior": polymer nanoparticle is in the interior of the BBB bilayer; "Outside": polymer nanoparticle has no contact with the BBB. In all the nanoparticle simulations, the elastic network was applied to preserve the solid-like structure.

b) Data of a control simulation of the BBB bilayer (*i.e.*, without nanoparticles).

**III. Supporting Figures**

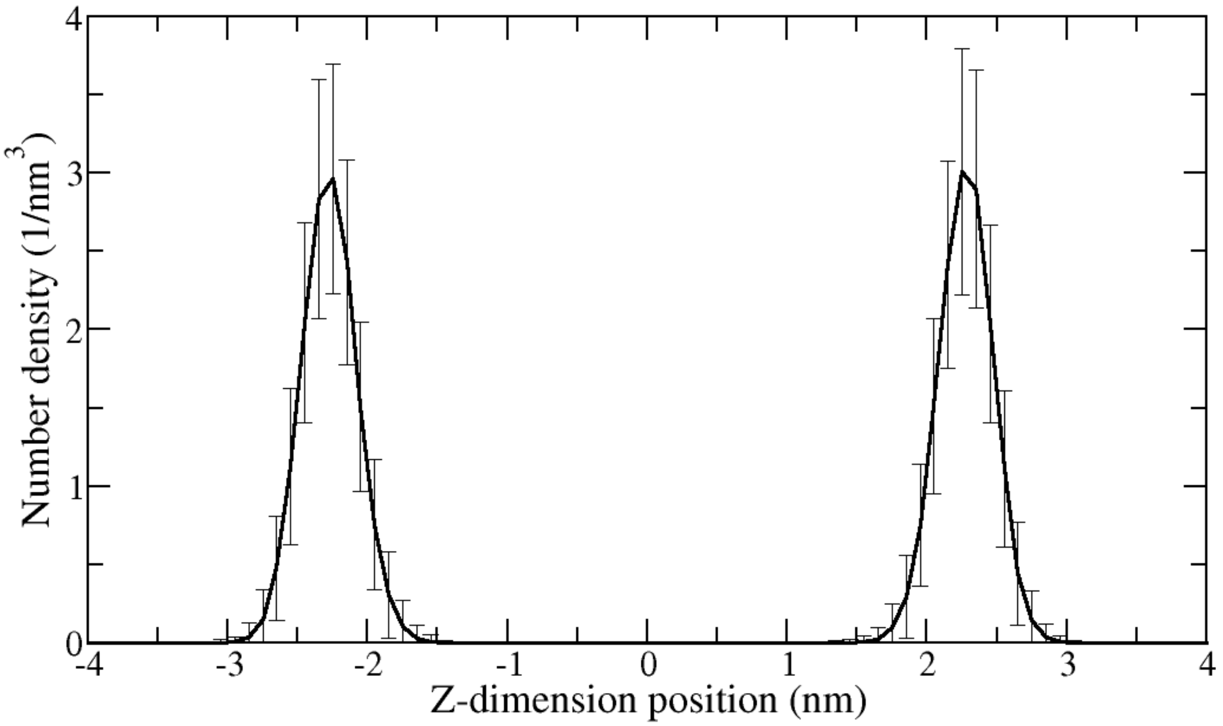


**Figure S1.** Number density profile of the lipid phosphorus atoms as a function of the distance to the BBB bilayer center in the control simulation (*i.e.*, in the absence of polymer nanoparticles). The error bars stand for the standard deviations calculated from the last 0.75 μs of the 0.8 μs control simulation. The thickness of the BBB bilayer is thus estimated to be around 4.5 nm.

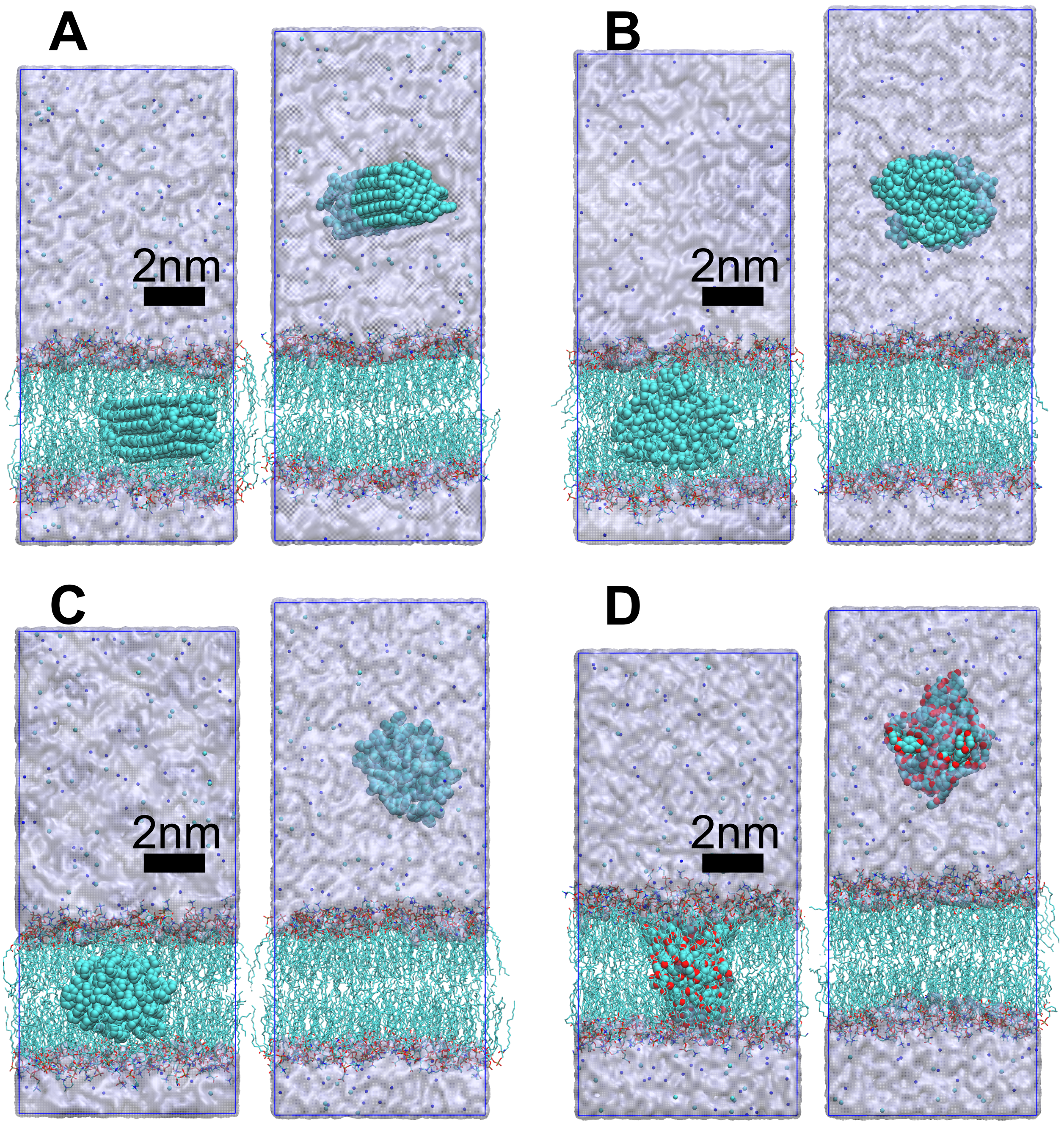


**Figure S2**. sMD simulation structures demonstrating (**A**) PE, (**B**) PP, (**C**) PS, and (**D**) PET nanoparticles located in (left) the BBB interior and (right) the water phase. The C/O/N/P/Na/Cl/H atoms are colored in cyan/red/blue/orange/white, respectively. $Na^{+}$ (blue) and $Cl^{-}$ (cyan) ions are also displayed. Polymer hydrogen atoms are omitted for the display. The scale bar is also presented.

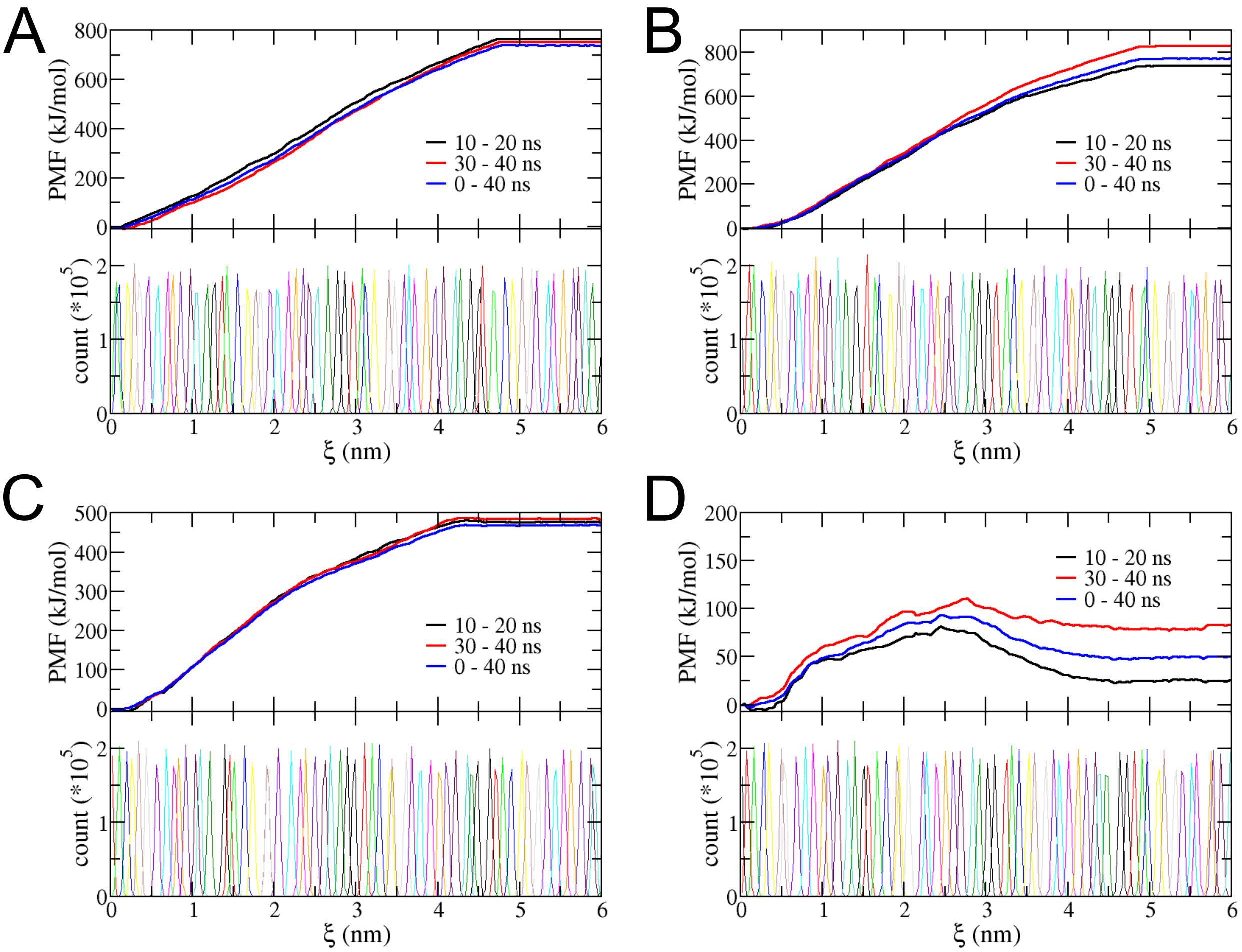


**Figure S3.** Convergence of the PMF calculations for the polymer nanoparticles. (**A**) PE, (**B**) PP, (**C**) PS, and (**D**) PET. In each subplot, presented in the top one are the PMF profiles calculated at different simulation periods, with the histogram distribution of the WHAM analysis provided in the bottom one.

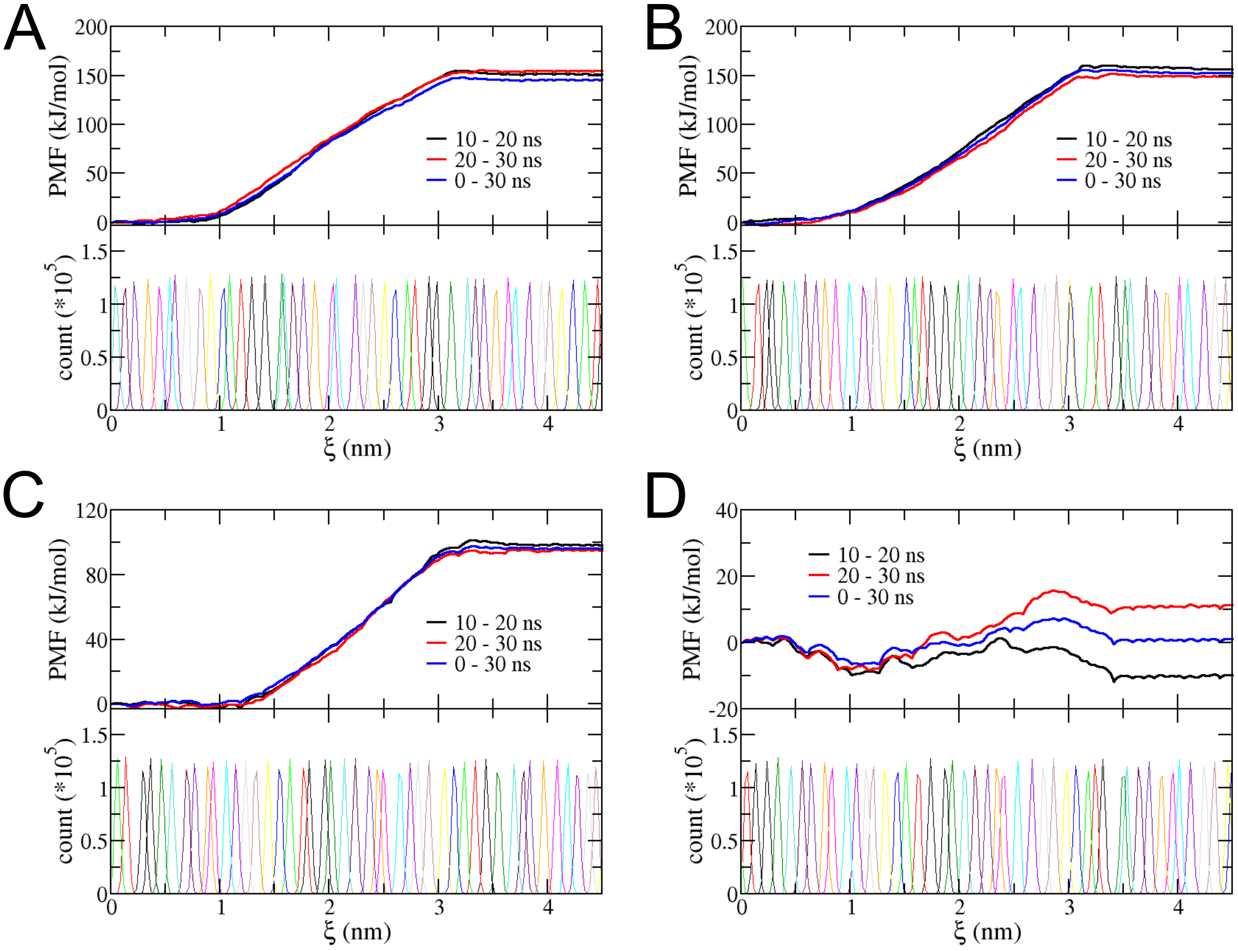


**Figure S4.** Convergence of the PMF calculations for single polymer chains. (**A**) PE, (**B**) PP, (**C**) PS, and (**D**) PET. In each subplot, presented in the top one are the PMF profiles calculated at different simulation periods, with the histogram distribution of the WHAM analysis provided in the bottom one.

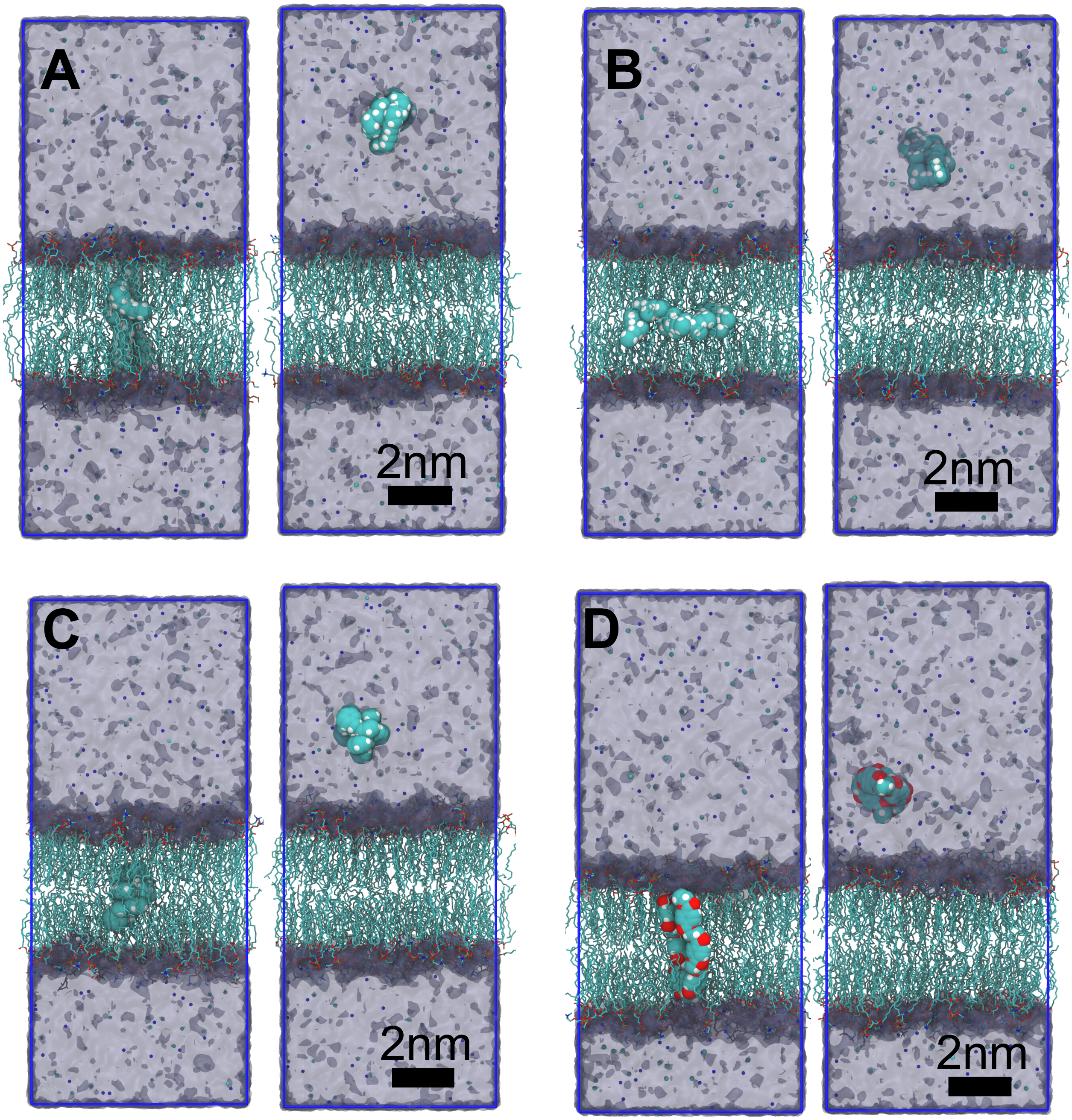


**Figure S5.** sMD simulation structures demonstrating single polymer chains of (**A**) PE, (**B**) PP, (**C**) PS, and (**D**) PET located in (left) the BBB interior and (right) the water phase. The C/O/N/P/Na/Cl/H atoms are colored in cyan/red/blue/orange/white, respectively. $Na^+$ (blue) and $Cl^-$ (cyan) ions are also displayed. The scale bar is also presented.
